# Stress-induced plasticity of a novel CRH^GABA^ projection disrupts reward behaviors

**DOI:** 10.1101/2022.07.01.498504

**Authors:** Matthew T. Birnie, Annabel K. Short, Gregory B. de Carvalho, Benjamin G. Gunn, Aidan L. Pham, Christy A. Itoga, Xiangmin Xu, Lulu Y. Chen, Stephen V. Mahler, Yuncai Chen, Tallie Z. Baram

## Abstract

Disrupted operations of the reward circuit are thought to underlie major emotional disorders including depression and drug abuse^1–3^. These disorders commonly arise following early life stress^4,5^; however, how stress early in life enduringly impacts reward circuit functions to promote disease remains unclear. Here, we discover and characterize a novel stress-sensitive reward-circuit projection connecting the basolateral amygdala (BLA) and nucleus accumbens (NAc) that co-expresses GABA and the stress-reactive neuropeptide corticotropin-releasing hormone (CRH). We then identify a crucial role for this projection in executing the disrupted reward behaviors provoked by early-life adversity (ELA): Chemogenetic and optogenetic stimulations of the CRH^GABA^ BLA→NAc projection in typically reared mice suppressed several reward seeking behaviors, recapitulating deficits resulting from ELA and demonstrating a key contribution of this pathway in the normal operations of the reward circuit. Next, inhibition of the CRH^GABA^ BLA→NAc projection in adult mice that experienced ELA restored typical reward behaviors in these mice, and, in contrast, had little effect in typically reared mice, indicating a selective ELA-induced maladaptive plasticity of this reward-circuit projection. We discover a novel, stress-sensitive, reward inhibiting projection from the BLA→NAc with unique molecular features, which may provide targets for intervention in disabling mental illnesses.

## Main

The brain is organized in circuits that orchestrate complex behaviors, including pleasure and reward^6^. The nucleus accumbens (NAc) is a major node of the reward circuit and is involved in pleasure and motivation processes^7^. Multiple afferents converge on the NAc to modulate reward behavior, including from the basolateral amygdala (BLA)^8,9^. The BLA mediates associative learning of aversive and appetitive stimuli, and glutamatergic BLA projections to the NAc facilitates reward behavior^10–12^.

The maturation of brain circuits during the early postnatal period is influenced by experiences including stress^13–16^. Stress experienced during critical early life periods (ELA) can exert enduring effects on brain circuits and their functions, resulting in mental health problems^17–19^. The mammalian brain is endowed with evolutionarily conserved stress-sensitive molecules, expressed in specific regions / brain-circuit nodes. Therefore, we reasoned that the effects of early-life stress on reward-circuit operation might take place by influencing the expression or function of neurons and projections expressing such stress-sensitive molecules. We focused on the neuropeptide CRH, an orchestrator of neuroendocrine stress signaling^20^ and an established modulator of NAc functions in a stress-dependent manner^21,22^. In addition, CRH-expression and function are enduringly influenced by early-life adversity (ELA) in several brain regions^23–25^. We thus mapped CRH-afferent projections to the NAc and describe here a novel, GABAergic CRH^+^ projection from the BLA→NAc and the use of chemogenetic and optogenetic strategies to determine its role in regulating reward behavior. We then establish that this projection mediates deficits in adult reward-seeking behaviors that are observed following ELA.

First, CRH-ires-CRE mice were injected with a CRE-dependent retrograde virus into the medial NAc shell (Fig. 1a, b) to retrogradely infect projection neurons^26,27^, including those originating in the BLA (Fig. 1c). The specificity of CRE-dependent viral expression to CRH^+^ neurons was confirmed with colocalization of endogenous CRH in viral-infected BLA cell bodies (Fig. 1d). To define the neuroanatomical distribution of the BLA-origin CRH^+^ projection in the NAc, we injected a CRE-dependent anterograde virus into the medial BLA (Fig. 1e-g,) and identified CRH^+^ BLA projection fibers and terminals in the medial NAc shell (Fig. 1h). A combination of viral reporter, fluorescent *in situ* hybridization and immunostaining identified this population of CRH^+^ neurons as GABAergic, rather than glutamatergic (Fig. 1i-k). Specifically, shown are images of BLA neurons from CRH ires-CRE mice injected with a CRE-dependent retrograde virus injected into the medial NAc shell to express tdTomato in BLA-origin cell bodies (Fig. 1i). These CRH^+^ neurons co-express two defining GABA markers: Gad67 (Fig. 1i) and the vesicular GABA transporter - vGAT (Fig. 1j). In contrast, they do not express a defining glutamatergic marker - CaMKII (Fig. 1k).

**Fig. 1:**
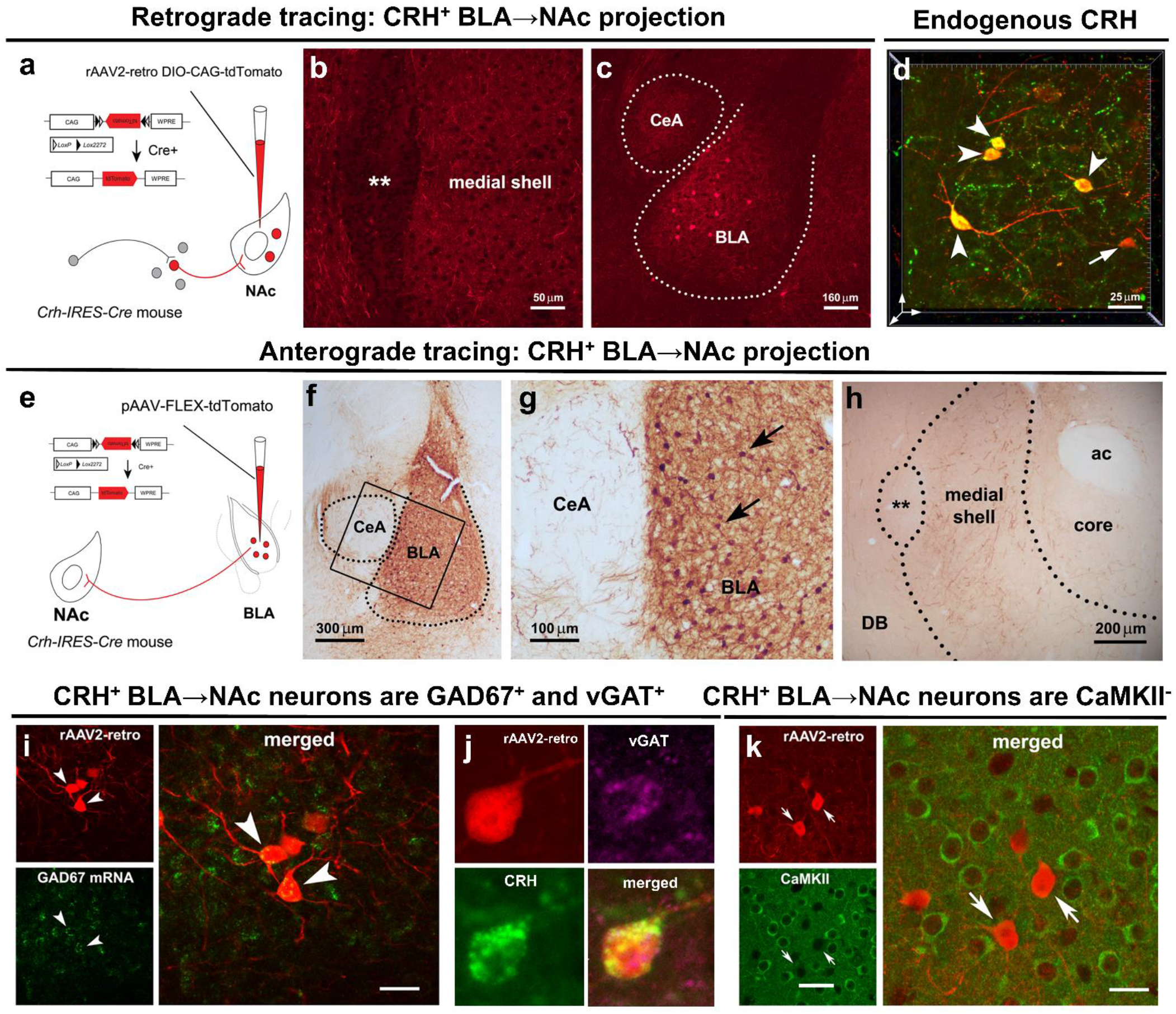
A novel projection of CRH^GABA^ neurons in the medial BLA to the medial NAc shell. **a**-**c**, Retrograde tracing of CRH^+^ neuronal inputs to medial NAc shell of CRH-ires-CRE mice. **a**, Schematic of construct and injection location of AAV2-retro-CAG-FLEX-tdTomato-WPRE virus that permits retrograde access to projection neurons providing afferent inputs to NAc. **b**, Example confocal micrograph of locally infected CRH^+^ axon terminals in medial NAc shell. **c**, Retrograde tracing identifies the medial BLA as a robust source of CRH^+^ NAc inputs. **d**, 3D image (z-stack; 0.5 μm steps) confirmed localization in the BLA of AAV-retro infected cells (red) that co-express endogenous CRH (green); dual labelled neurons = yellow. **e**-**g**, Anterograde tracing of CRH^+^ axonal projections from BLA to medial NAc shell. **e**, The AAV1-DIO-tdTomato construct and the viral genetic experimental design. **f**, Virus injection is confined to the BLA, **g**, and absent from the central amygdala, shown by selective expression of tdTomato in BLA CRH^+^ neurons. **h**, BLA-origin CRH^+^ axons and terminals in the medial NAc shell. **i**-**k**, Virus injection into the medial NAc shell retrogradely infected somata in the BLA. **i**, Combined fluorescence *in situ* hybridization (FISH) and immunostaining with GAD67 mRNA in CRH^+^ cells in the BLA. Arrowheads point to co-localized GAD67 mRNA and virus-reporter labelling. **j**, a BLA→NAc cell (red) co-expresses endogenous CRH (green) and vGAT (magenta), but **k**, does not co-express the glutamatergic marker CaMKII. ** = Major Island of Calleja, ac = anterior commissure. Scale bars in **i** and **k** = 10μm; **j** = 50μm.

To substantiate that the CRH^+^ BLA→NAc projection evokes inhibitory currents, we performed whole cell patch clamp recordings in the medial NAc shell of CRH-ires-CRE mice injected with a CRE-dependent ChR2-EYFP in the BLA (Fig. 2a). Optical stimulation of CRH^+^ BLA-origin fibers in the NAc reliably evoked inhibitory postsynaptic currents (oIPSCs) (Fig. 2b) in medium spiny neurons (MSNs) (Fig. 2c). The optically evoked inhibitory postsynaptic responses were consistently blocked by superfusion with the GABA_A_ receptor antagonist, picrotoxin (100 μM) (Fig. 2d, e). In addition, in a confirmed oIPSC recorded cell, optical stimulation did not yield excitatory postsynaptic currents (oEPSCs) (Fig. 2f), even though spontaneous EPSCs were apparent (Fig. 2g). Together, these data identify a novel, inhibitory CRH^GABA^ BLA→NAc projection that is distinct from the well-described glutamatergic BLA→NAc projection^10,11,28^.

**Fig. 2:**
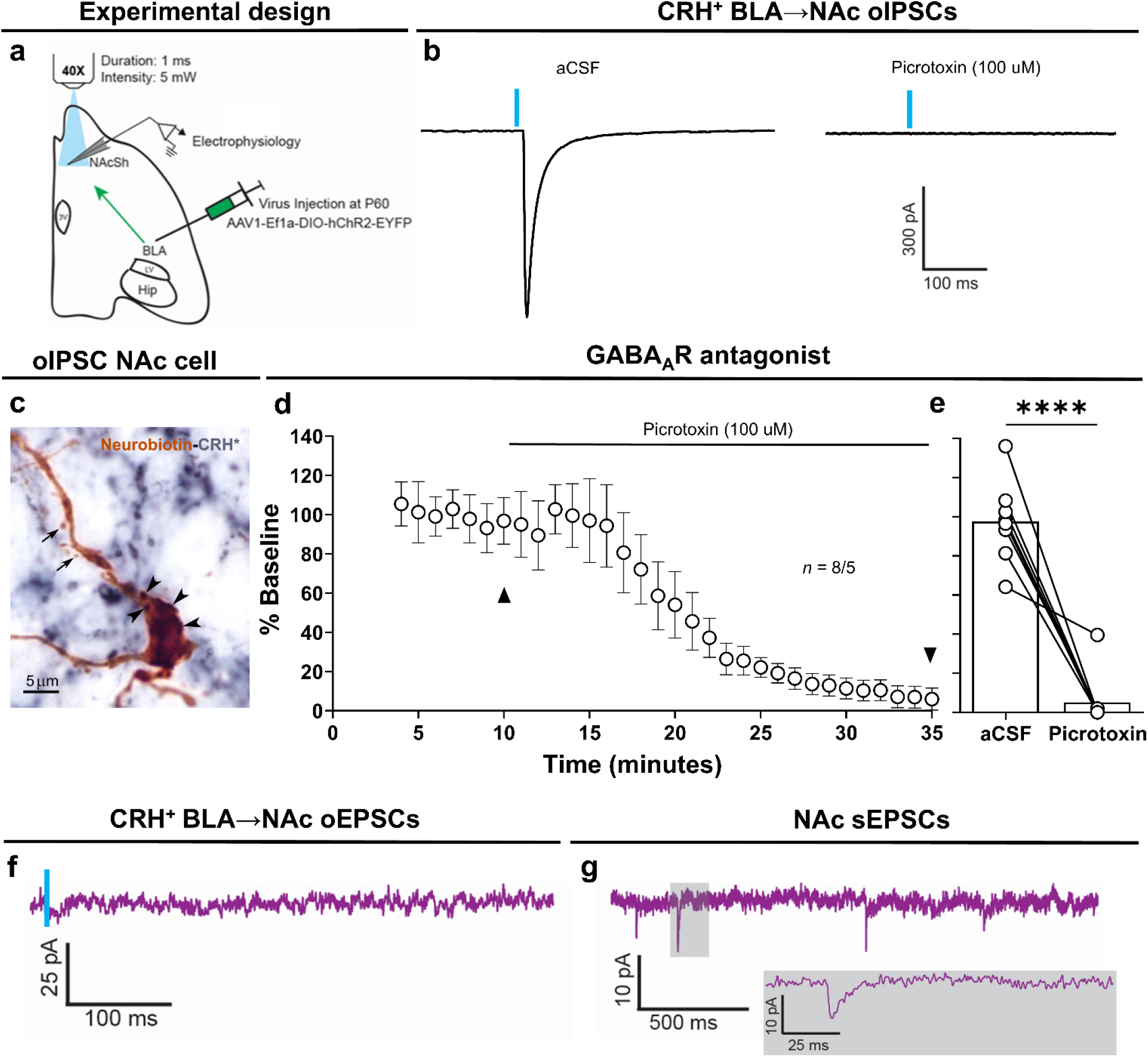
Optical stimulation of CRH^+^ BLA-origin axons in the NAc evokes exclusively IPSCs. **a**, Schematic of experimental design for electrophysiology recordings in the whole-cell patch-clamp configuration. Horizontal brain slices containing BLA and NAc from CRH-ires-CRE mice that were injected with Cre-dependent ChR2-EFYP in BLA. **b**, Representative traces of optically evoked IPSCs (oIPSCs). These were blocked in the presence of GABA_A_ receptor antagonist picrotoxin. **c**, A neurobiotin (brown) filled neuron from which oIPSCs were recorded. Note spines (arrows) suggesting the cell is a medium spiny neuron (MSN). Arrowheads denote ChR2-expressing boutons from BLA-origin CRH^+^ axons (CRH; blue) on the soma. **d**, Time-course plot of oIPSCs amplitudes throughout the recording and following application of picrotoxin. **e**, oIPSCs amplitudes pre and post picrotoxin (*n* = 8 neurons, 5 mice). Black triangles in **d** denote trace recordings in **b** and timepoint analysis in **e. f**, Representative trace of a NAc cell showing no response from optically evoked EPSC (oEPSC) at -70 mV, obtained after verifying oIPSCs at 0 mV in the presence of picrotoxin. **g**, Representative trace showing spontaneous EPSCs (sEPSCs) were still present; grey box shows magnified view recording. In **d**, circles represent mean ± SEM. Paired *t*-test in **e. e**, oIPSCs pre vs post PTX: *P* < 0.0001. In all figures, \**P* < 0.05, \*\**P* < 0.01, \*\*\**P* < 0.001 and \*\*\*\**P* < 0.0001. NS, not significant.

To investigate the function of this CRH^GABA^ BLA→NAc projection, we targeted it during reward behaviors using chemogenetic and optogenetic strategies. Specifically, in typically reared mice, we microinjected CNO^29,30^ into the medial NAc shell (Ext. Fig. 3a) to stimulate the CRH^GABA^ BLA-origin fibers in the NAc, during two different reward tasks – one centered on palatable food consumption and the other involving sex cues^31^. Bilateral microinfusion of CNO (1 mM) into the medial NAc shell reduced palatable food consumption (Fig. 3b). This suppression was also evident in the sex-cue task (Fig. 3c). Notably, these results were not a result of off-target effects of CNO (Ext. Fig. 3e), as non-reward behaviors were unaffected, and we confirmed the specific activation of hM3Dq-expressing CRH^+^ neurons by CNO via selective Fos expression in the BLA 90 minutes following injection (Ext. Fig. 3c, d). We then used a second, independent technology, optogenetics, injecting a CRE-dependent ChR2-EYFP in the BLA bilaterally of CRH ires-CRE mice, and implanting bilateral optic fibers directly above the medial NAc shell to optically stimulate projection fibers (Ext. Fig. 3b). Blue light (473 nm) stimulation (10Hz) suppressed palatable food consumption (Fig. 3d) and the preference for a sex-cue (Fig. 3e). Again, potential off-target effects of virus and/or light stimulation did not contribute to these outcomes (Ext. Fig. 3f-i). Collectively, these findings describe an unexpected reward-suppression role for this CRH^GABA^ BLA→NAc projection, which differs from the well-established, reward enhancing glutamatergic projection connecting the BLA and NAc^10,11^. Crucially, CRH expression and function are often modulated by stress^24,32^, therefore these data supported the possibility that this projection might mediate suppressed reward behaviors often observed after adult or developmental stresses.

**Fig. 3:**
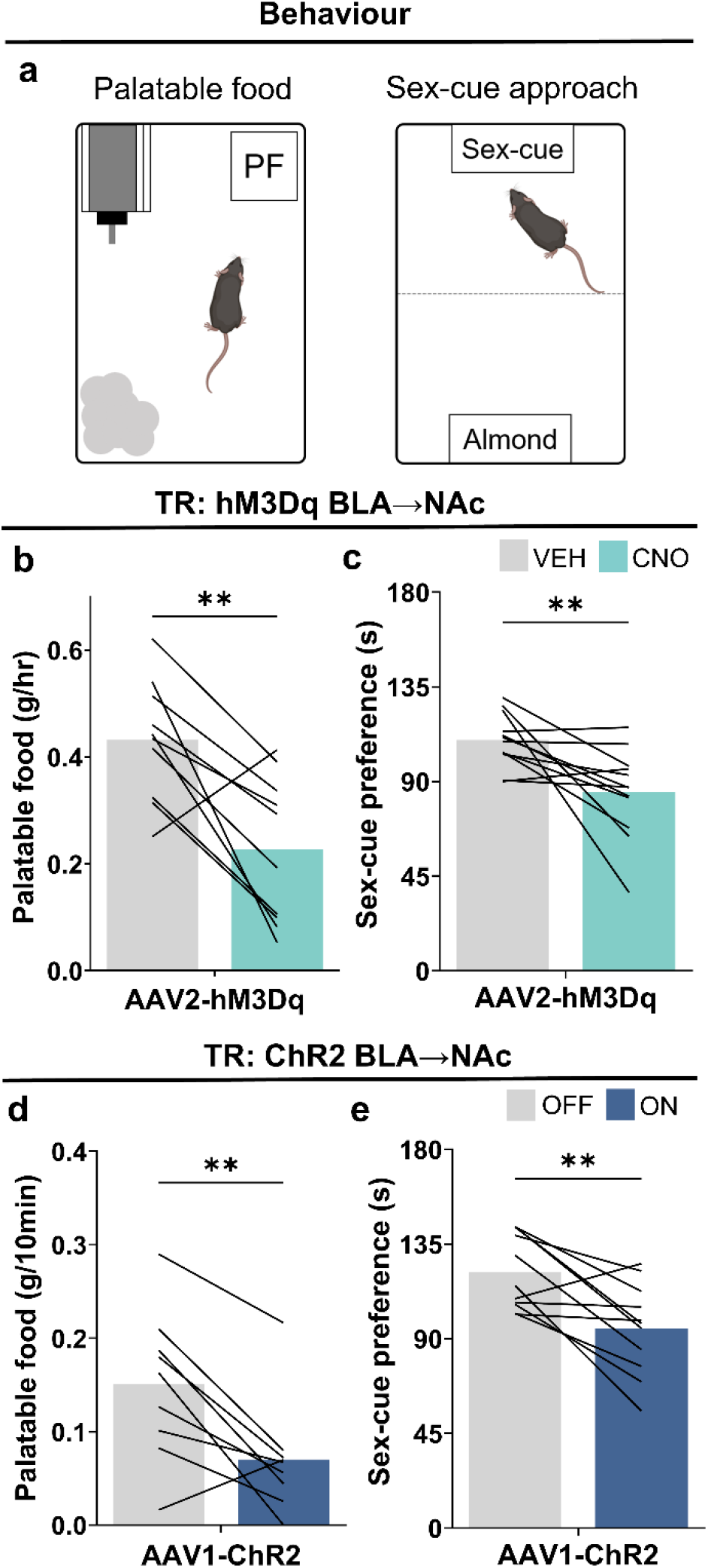
Stimulating the CRH^GABA^ BLA→NAc projection suppresses reward in typically reared mice. **a**, Schematic of reward tasks. **b, c**, Stimulating the CRH^GABA^ BLA→NAc projection with microinfusion of CNO in the medial NAc shell reduced **b**, palatable food consumption (*n* = 10 mice) and **c**, approach time to a sex-cue (*n* = 12 mice). **d, e**, Stimulating the CRH^GABA^ ChR2-expressing BLA→NAc projection decreased D, palatable food consumption (*n* = 9 mice) and **e**, approach time to a sex-cue (*n* = 11 mice). In **b**-**e**, bars represent mean. Paired *t*-tests (**b**-**e**). **b**, hM3Dq BLA→NAc: *P* = 0.0037; **c**, hM3Dq BLA→NAc: *P* = 0.0096; **d**, ChR2 BLA→NAc: *P* = 0.0064; **e**, ChR2 BLA→NAc: *P* = 0.0034.

A large body of work documents a disruption of reward behaviors following ELA in both humans and experimental models^33–35^. To determine whether the CRH^GABA^ BLA-NAc projection contributes to these deleterious effects, we first determined consequences of ELA on reward behaviors. We used a well-established model of ELA during a sensitive developmental period^36,37^ (Fig. 4a), in which poverty is simulated via a resource-scarce environment during postnatal days 2-10 in mice^17^. In adult ELA mice, preference for palatable food, but not regular chow, was diminished (Fig. 4b). Similarly, preference for a sex-cue (Fig. 4c) and sucrose (Fig. 4d) were also reduced, whereas effect of ELA on the open field or swim stress tasks were not apparent (Ext. Fig. 4a, b). Strikingly, the reward deficits in adult ELA mice recapitulated the effects of chemogenetic and optogenetic stimulation of the CRH^GABA^ BLA→NAc projection in typically reared (TR) mice (Fig. 3), identifying a possible role for this projection in ELA-induced reward deficits^14^. To test this, we suppressed the activity of the CRH^GABA^ BLA→NAc projection during reward tasks in adult ELA mice. Specifically, we provided CRH-ires-Cre ELA mice with bilateral injections of a CRE-dependent inhibitory DREADD (hM4Di) into the BLA and then infused CNO (1 mM) directly into the medial NAc shell (Fig. 4e, f). Inhibition of the BLA-origin fibers in the NAc rescued the consumption of palatable food in adult ELA-experienced mice (Fig. 4e) and restored the preference for a sex-cue (Fig. 4f). This approach did not alter locomotion or induce an anxiety-like phenotype (Ext. Fig. 4c, d). Crucially, inhibiting this projection in TR mice did not increase palatable food consumption (Fig. 4g) or the preference for a sex cue (Fig. 4h). Together, these data suggest that inhibiting the CRH^GABA^ BLA→NAc projection in ELA, but not TR mice, increases reward behavior, indicating a selective ELA-induced maladaptive plasticity of this reward-circuit projection.

**Fig. 4:**
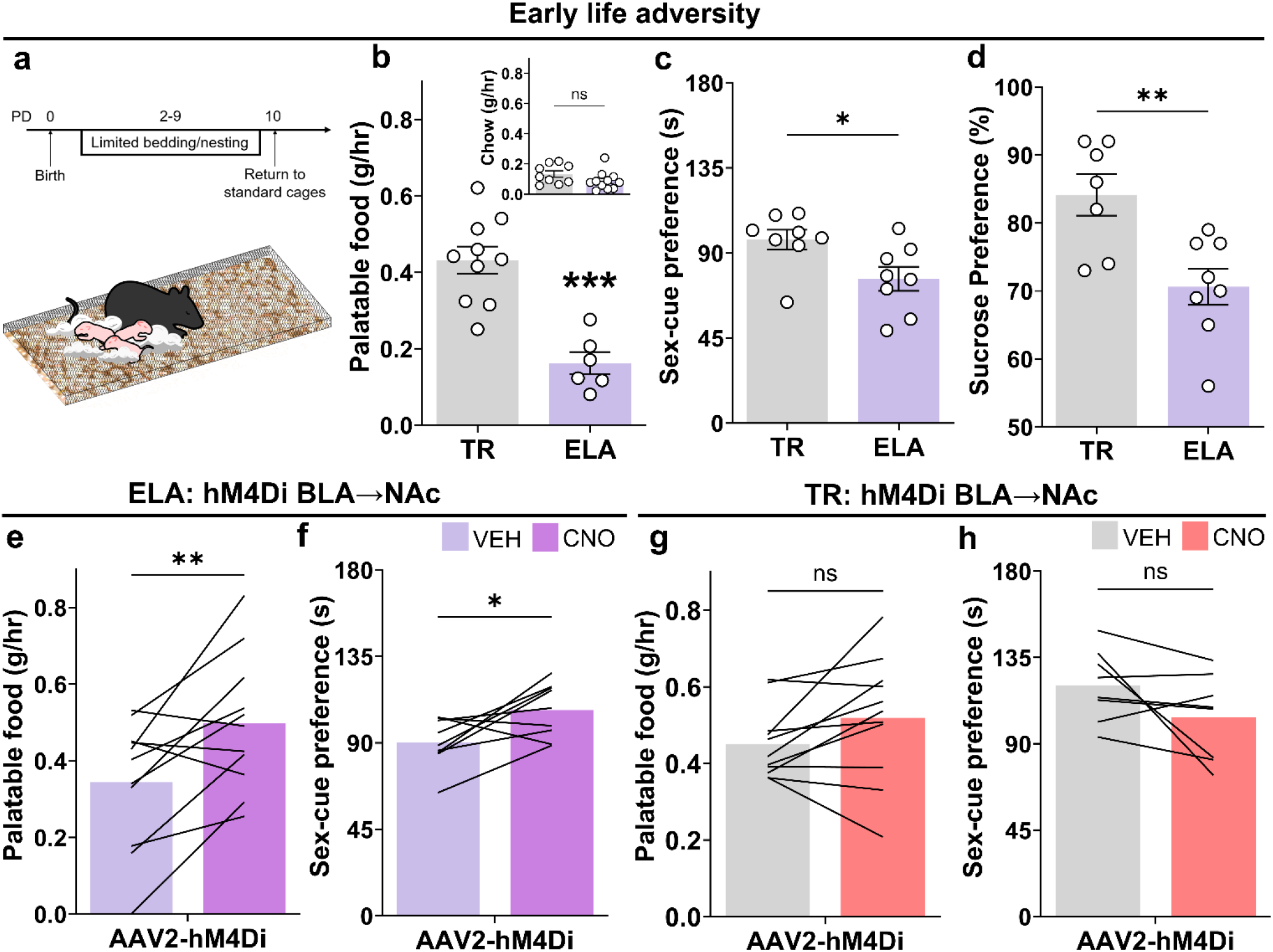
Inhibiting the CRH^GABA^ BLA→NAc projection rescues reward deficits following early life adversity. **a**, Timeline and environment (limited bedding and nesting) of ELA. **b**-**d**, ELA reduced palatable food consumption (*n* = 16 mice; *TR* = 10, *ELA* = 6) yet did not alter regular chow consumption (*n* = 20 mice; *TR* = 9, *ELA* = 11). ELA reduced preference for a sex-cue (*n* = 16 mice; *TR* = 8, *ELA* = 8) and preference for sucrose (*n* = 15 mice; *TR* = 7, *ELA* = 8). **e**, Inhibiting the CRH^GABA^ BLA→NAc projection with microinfusion of CNO in the medial NAc shell increased palatable food consumption (*n* = 11 mice) and **f**, preference for a sex-cue (*n* = 9 mice). Inhibiting the CRH^GABA^ BLA→NAc projection in TR mice did not increase **g**, palatable food consumption (*n* = 11 mice) or **h**, preference for a sex-cue (*n* = 8 mice). In **b**-**d**, bars represent mean ± SEM; in **e**–**h**, bars represent mean. Unpaired *t*-tests (**b, d**), unpaired *u*-test (**c**), paired t-tests (**e**-**h**). **b**, TR vs. ELA main: *P* = 0.0001, inset: *P* = 0.1263; **c**, TR vs. ELA: *P* = 0.0281; **d**, TR vs. ELA: *P* = 0.0053; **e**, ELA hM4Di BLA→NAc: *P* = 0.0037; **f**, ELA hM4Di BLA→NAc: *P* = 0.0246; **g**, TR hM4Di BLA→NAc: *P* = 0.1036; **h**, TR hM4Di BLA→NAc: *P* = 0.1139.

Exposure to severe or chronic stressors during sensitive periods in development can produce profound and often lasting changes to brain operations that can be detrimental to mental health^18,38^. A majority of the world’s adults have experienced a form of ELA, which precedes--and likely contributes to--affective disorders including depression, drug use and obsessive risk-taking^17,39^. These major public health challenges are associated with aberrant reward processing^40,41^. Remarkably, the cellular and circuit foundations of ELA-associated disruption of reward-related behaviors have remained largely elusive^17^. In this study, we first discovered an inhibitory BLA→NAc projection responsible for ELA-provoked deficits in reward behavior. Unlike the canonical glutamatergic projection from BLA→NAc^10,11^, this novel stress-sensitive CRH^GABA^ BLA→NAc projection suppresses several types of reward behaviors in typically reared mice. These findings highlight the emerging complexity of reward circuit operations, afforded by discoveries of novel cell-type specific projections that anatomically parallel established connections (e.g., BLA to NAc), yet exert different and sometimes opposing effects on behavior. This work also identifies the BLA as a novel locus of neurons expressing CRH, a peptide known to be released during stress in several brain regions^21,22,32,42^. Most studies on amygdalar CRH have centered on the central nucleus^43– 46^, which is rich in CRH-expressing cells yet does not project directly to the NAc^27^ (Fig. 1). We delineate a substantial population of CRH^GABA^ neurons in the BLA which project monosynaptically to the NAc to influence reward behavior, and importantly, whose function is impacted by ELA, promoting pathological deficits in these behaviors. Advancing our knowledge of stress-mediated control of reward behavior takes us a step closer to understanding the mechanisms of several prevalent major mental illnesses and provide novel targets for the prevention and intervention of such stress-related disorders.

---

**Ext. Fig. 3:**
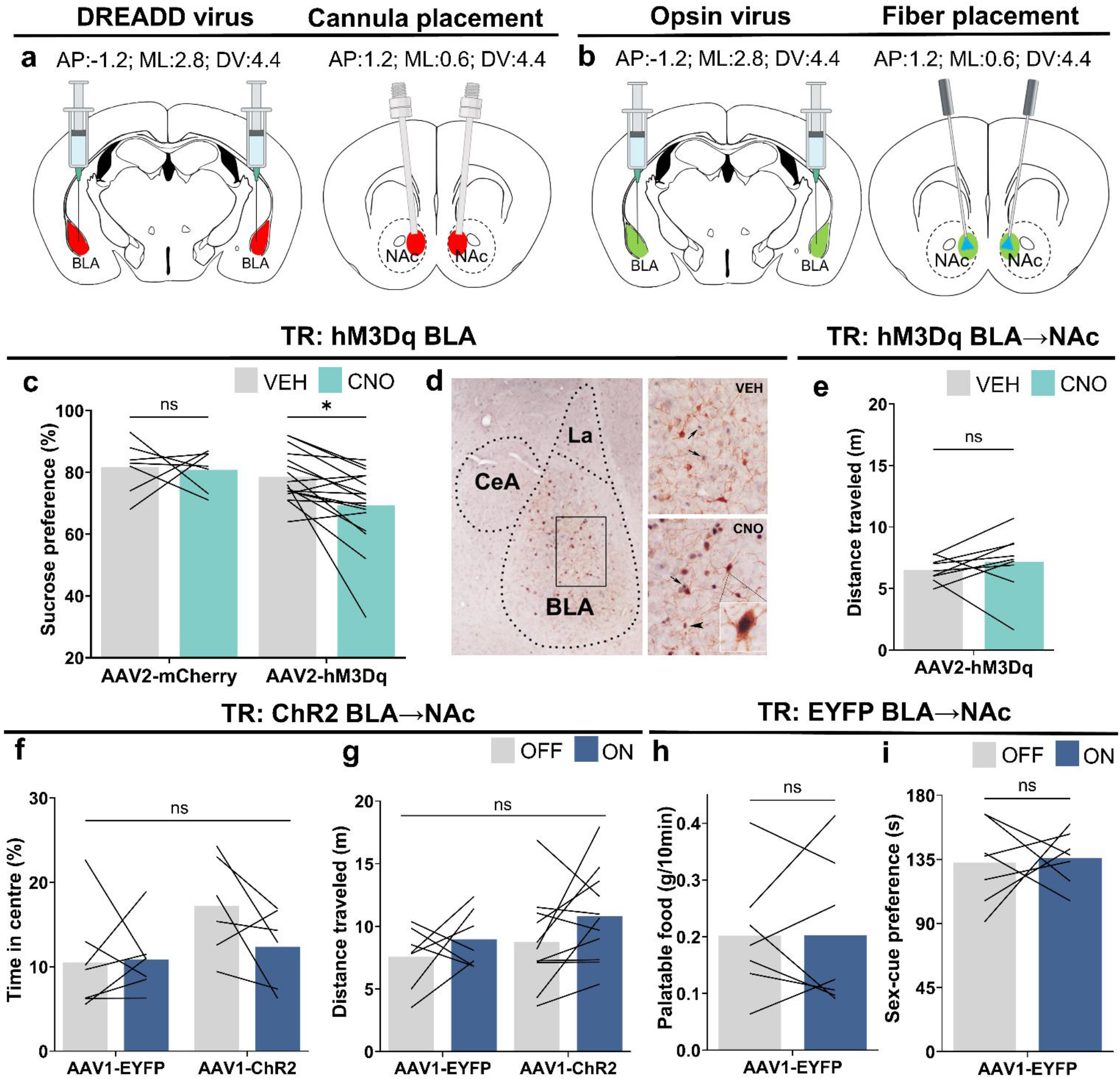
Stimulating the CRH^GABA^ BLA→NAc projection suppresses reward. Coordinate locations of **a**, DREADD injection and CNO infusion and **b**, opsin injection and fiber placement. **c**, Stimulating CRH^GABA^ AAV2-hM3Dq-expressing cells in the BLA decreased sucrose preference with systemic injection (*n* = 23 mice; *mCherry* = 7, *hM3Dq* = 16). **d**, Example Fos expression in CRH^GABA^ AAV2-hM3Dq-expressing BLA neurons 90 min post injection of vehicle or CNO. **e**, Microinjections of CNO in the medial NAc shell of mice expressing AAV-DIO-mCherry to target the CRH^+^ BLA-origin projection did not affect distance traveled during the sex-cue task (*n* = 9 mice). **f, g**, Optical stimulation of the CRH^+^ BLA→NAc projection did not alter **f**, time spent in center in an open field task (*n* = 13 mice; *EYFP* = 7, *ChR2* = 6) or **g**, distance travelled during the sex-cue task (*n* = 18 mice; *EYFP* = 7, *ChR2* = 11). **h, i**, Optical stimulation of the BLA→NAc projection did not affect **h**, consumption of palatable food (*n* = 7 mice), or **i**, sex-cue preference (*n* =7 mice). In **c, e**–**i**, bars represent mean. Paired *t*-tests (**e, h, i**), Two-way ANOVA with repeated measures followed by Sidak’s post hoc test (**c**), Two-way ANOVA with repeated measures (**f, g**). **c**, hM3Dq BLA: *F* = 6.075, DFn = 1, DFd = 42, *P* = 0.0179; **e**, hM3Dq BLA→NAc: *P* = 0.4523; **f**, ChR2 BLA→NAc: *F* = 4.083, DFn = 1, DFd = 22, *P* = 0.0557; **g**, ChR2 BLA→NAc: *F* = 1.827, DFn = 1, DFd = 32, *P* = 0.186; **h**, EYFP BLA→NAc: *P* = 0.9923; **i**, EYFP BLA→NAc: *P* = 0.8344.

**Ext. Fig. 4:**
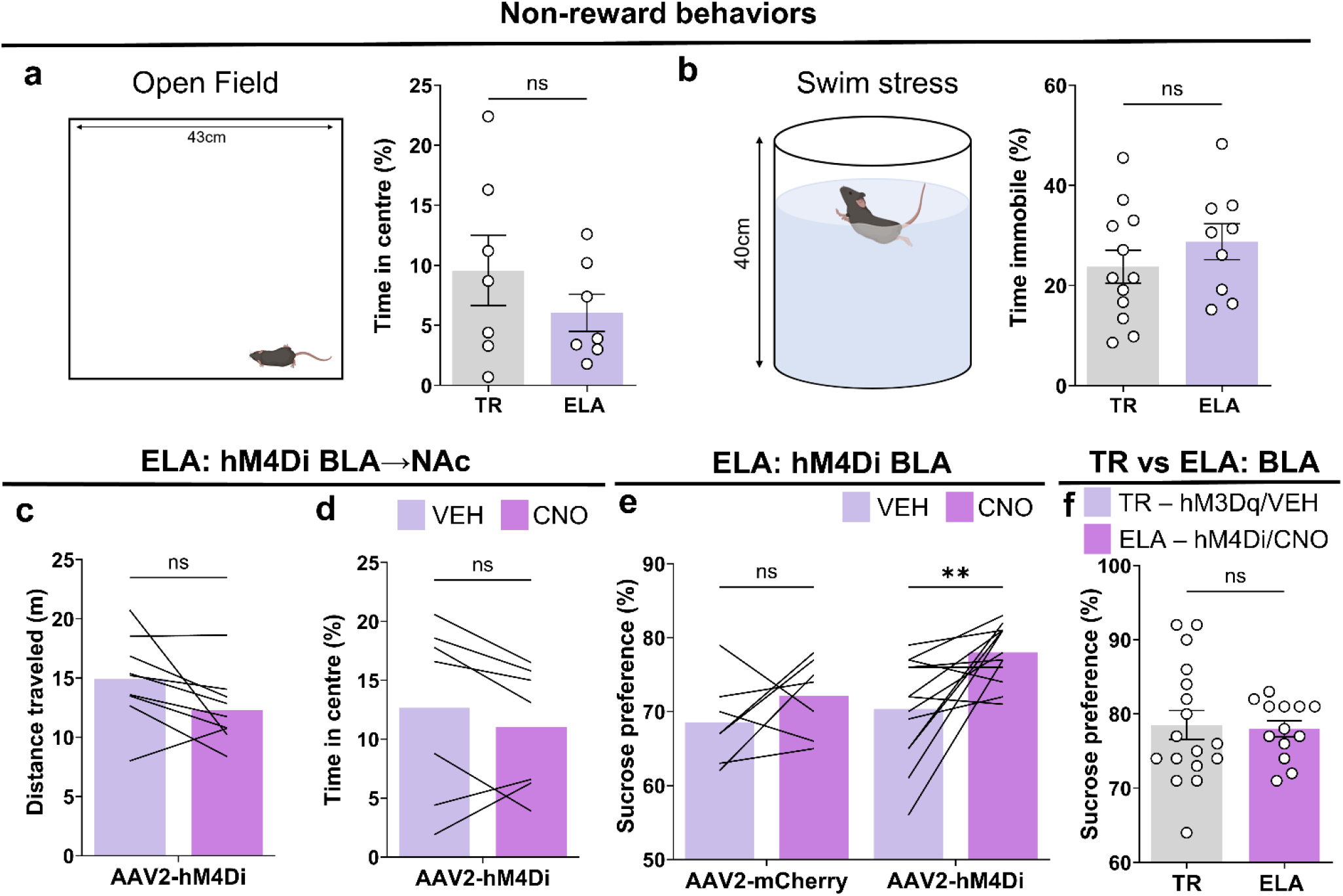
Inhibiting the CRH^GABA^ BLA→NAc projection rescues reward deficits following early life adversity. **a, b**, ELA mice do not have altered response to stress-mediating, non-reward tasks, for example **a**, time spent in the center in an open field (*n* = 14 mice; *TR* = 7, *ELA* = 7) or **b**, immobility time during swim stress (*n* = 21 mice; *TR* = 12, *ELA* = 9). **c**, Inhibiting the CRH^GABA^ BLA→NAc projection did not affect distance traveled during the sex-cue task (*n* = 9 mice) or **d**, time spent in the center in an open field task (*n* = 7 mice). **e**, Inhibiting CRH^GABA^ BLA neurons enhances preference for sucrose in ELA mice (*n* = 20 mice; *mCherry* = 7, *hM4Di* = 13), to levels seen in **f**, TR mice (*n* = 30 mice; *mCherry* = 17, *hM4Di* = 13. In **a, b, f**, bars represent mean ± SEM. In **c**-**e**, bars represent mean. Unpaired t-tests (**a, b, f**), paired t-tests (**c, d**), Two-way ANOVA with repeated measures followed by Sidak’s post hoc test (**e**). **a**, TR vs ELA: *P* = 0.3083; **b**, TR vs ELA: *P* = 0.3258; **c**, BLA→NAc hM4Di: *P* = 0.0609; **d**, BLA→NAc hM4Di: P=0.2748; **e**, ELA BLA: *F* = 4.25, DFn = 1, DFd = 36, *P* = 0.0465. **f**, BLA TR (hM3Dq-Veh) vs ELA (hM4Di-CNO): *P* = 0.8290.

## MATERIALS AND METHODS

### Experimental animals

Male B6(Cg)-Crh^tm1(cre)Zjh^/J (CRH-ires-CRE) mice were used in all experimental procedures. Food and water were available ad libitum. Mice were bred in house from Jackson Laboratories stock (Stock no: 012704). All mice were housed in standard conditions at 72°F and 42% humidity and a 12-hour light-dark cycle (Lights ON: 0600; Lights OFF: 1800). All mice were single housed following surgical procedures for experimental purposes. All experimental procedures were approved by the University of California-Irvine Institutional Animal Care and Use Committee (AUP 21-128) and were in accordance with the guidelines from the National Institute of Health.

### Limited bedding and nesting paradigm

Early life adversity was induced on postnatal day 2 to 9 using our laboratory’s standard protocol, as described previously^30^. Briefly, routine plastic mouse cages (L37 x W20 x H17 cm) were fitted with an aluminum mesh platform sitting ∼2.5cm above the cage floor. Bedding was reduced to cover the cage floor sparsely, and one-half of a single nestlet was provided for nesting material. Typically reared (TR) dams and pups resided in standard bedding cages, but were moved to new cages on P2 to control for handling during the LBN setup. TR and LBN cages were undisturbed from P2-P9, and housed in a quiet room. On P10, both TR and ELA groups were transferred to fresh, routine cages.

### Animal surgery

Mice were anaesthetized with either ketamine/xylazine (160/12 mg/kg body weight) or isoflurane (1-1.5%) for stereotaxic surgery. For retrograde tracing experiments, mice received a unilateral injection (0.2ul, 20-30nL/min) of AAV2-retro-CAG-FLEX-tdTomato-WPRE into the left medial NAc shell (AP: 1.18, ML: 0.6, DV: 4.55). For anterograde tracing experiments, mice received a unilateral injection (0.2-0.3ul) of AAV1-DIO-tdTomato into left hemisphere BLA (AP: -1.35, ML: +/-2.75, DV: 4.5). For in vivo chemogenetic excitation during behavior testing, mice were injected with the AAV2-hM3D(Gq)-mCherry virus or the same viral vectors carrying mCherry alone (5×10^12^ vector genomes/mL) bilaterally into the BLA. For in vivo chemogenetic inhibition during behavior testing, mice were injected with the AAV2-hM4D(Gi)- mCherry bilaterally into the BLA. For intra NAc drug infusion, guide cannulae were implanted over the medial NAc shell (AP: 1.18, ML: +/-0.6, DV: 4.2). For in vivo optogenetic excitation during behavior, and in vitro electrophysiology experiments, mice were injected with the AAV1-EF1a-DIO-hChR2(h134R)- EYFP virus or the same viral vector carrying EYFP alone, bilaterally into the BLA. For projection specific targeting during behavior testing, optic fibers (0.22 NA, 100μm diameter) were placed dorsal to EYFP-expressing axonal terminals in the medial NAc shell (AP:1.18, ML: +/-0.6, DV:4.3).

### Chemogenetic studies

For excitatory (AAV2-hM3D(Gq)-mCherry) and inhibitory (AAV2-hM4D(Gi)-mCherry) DREADD experiments. For intraperitoneal injection of 1mg/kg Clozapine-N-Oxide (CNO) (NIMH, # MH-929, Batch #14073-1), CNO was dissolved in 1% Dimethyl sulfoxide (DMSO) in phosphate buffer saline (PBS). For microinfusions of CNO (CNO dihydrochloride, HelloBio #HB6149, Batch #E1169-1-2) into the medial NAc shell, 1mM CNO (0.2ul) in artificial cerebral spinal fluid (in mM: 124 NaCl, 3 KCl, 833 26 NaHCO3, 1.4 NaH2PO4, 1 MgSO4, 10 D-Glucose, 2 CaCl2).

### Optogenetic studies

For in vivo experiments, <5mW of blue light (95-159 mW/mm^2^ at the tip) was generated by 473-nm μLED (Prizmatix, US), and bilaterally delivered through two fiber optic patch cords (0.22 NA, 200um diameter, Prizmatix, US). Light delivery was controlled using a pulse generator (PulserPlus, Prizmatix, US) to deliver 5ms light pulse trains at 10 Hz^11^ during behavior testing.

For ex vivo electrophysiology recordings, <5mW of blue light was generated by a 473-nm LED (Sola Light Engine, Lumencor, US) and delivered through a 40X objective. Frequency (0.1Hz) and pulse duration (1ms) were controlled by LLE Sola-SE2 software (Lumencor, US).

### Electrophysiology

Slice Preparation: CRH-Cre mice (P60) injected with AAV1-Ef1a-DIO-hChR2-EYFP (Addgene cat # 20298-AAV1) in the basolateral amygdala (BLA) were deeply anesthetized with isoflurane and quickly decapitated (P120). Acute horizontal slices (300 μm) containing both BLA and nucleus accumbens (NAc) were obtained using a vibratome (Leica V1200S) in ice-cold N-Methyl D-Glucamine (NMDG) cutting solution: containing (in mM): 110 NMDG, 20 HEPES, 25 glucose, 30 NaHCO31.2 NaH2PO4, 2.5 KCl, 5 sodium ascorbate, 3 sodium pyruvate, 2 Thiourea, 10 MgSO4-7 H2O, 0.5 CaCl2, 305-310 mOsm, pH 7.4. Slices equilibrated in a homemade chamber for 25 – 30 minutes (31° C) and an additional 45 minutes in room temperature aCSF containing (in mM): 119 NaCl, 26 NaHCO3, 1 NaH2PO4, 2.5 KCl, 11 Glucose, 10 Sucrose, 2.5 MgSO4-7 H2O, and 2.5 CaCl2 (pH 7.4) before being transferred to a recording chamber. Recording solution was the same as equilibration solution only with 1.3 mM MgSO4-7 H2O. All solutions were continuously bubbled with 95% O2/5%CO2.

Whole-Cell Patch-Clamp: Whole-cell patch clamp recordings were obtained in the medial shell of the NAc (NAcSh). Data were collected with a Multiclamp 700B amplifier, Digidata 1550B (Molecular devices), and using Clampex 11 (pClamp; Molecular Devices, San Jose, CA). All recordings were acquired in voltage clamp at 34° Celsius and were digitized at 10 kHz and low pass filtered at 2 kHz. Patch pipette was filled with internal solution containing (in mM): 143 CsCl, 10 HEPES, 0.25 EGTA, 5 Phosphocreatine, 4 MgATP, 0.3 NaGTP (295 – 305 mOsm, pH 7.4 with CsOH) and 1mg/ml NeuroBiotin. CNQX (50μM) and AP5 (10μM) were added to aCSF to block glutamate receptors. All pipettes (3 - 4 MΩ) were pulled from borosilicate glass (Narishige PC-100). Series resistance (Rs) was monitored throughout the recording for patch sealing. Once whole cell configuration was obtained, holding potential was set at -70 mV and a 1 ms, 5 mW light pulse was delivered every 10 seconds (0.1 Hz) through a 40X objective. Once current response was obtained, picrotoxin (100 μM) was washed in the recording chamber to verify that optically evoked inhibitory postsynaptic current (oIPSC) was mediated by GABAa receptor activation. Once recording was complete slices were transferred to 4% PFA overnight at 4° Celsius then to 0.1 M PB for post-hoc processing of NeuroBiotin.

### Behavioural assays

All training and testing took place in standard housing mouse cages (34 × 18 cm), apart from open field (45 × 45 cm) and swim stress. All behavior was carried out in low light (<15 lux) and during the animal’s active period. Prior to testing start, all mice were placed in the behavior room for at least 1 hr to acclimatize to their surroundings.

#### Sucrose Preference Task

Single housed mice were habituated to the presence of two drinking bottles (both containing water) in their home cage for 3 days. Following acclimatization, one of the bottles was replaced with 1% sucrose solution and intake was measured 4 hours after IP CNO injection for 3 consecutive days during the early active period. The bottles switched sides daily to prevent side bias. Total consumption of water and sucrose solution was measured at the end of each time session by weighing the bottles. Sucrose preference was defined as the ratio of the consumption of sucrose solution vs the consumption of both water and sucrose solution combined. Testing was performed over 3 consecutive days. For chemogenetic experiments, mice were injected 1 hour prior to the start of the 4 hour test session.

#### Sex Cue Approach

Urine was collected from females in the estrus phase of the estrous cycle. 60ul of estrus urine and 60ul of a non-reward scent (Almond; Pure Almond extract, Target) were pipetted onto cotton tip applicators and fixed to the home cage of single housed mice at either end. The mouse was given free access to both tips for 3 minutes. For chemogenetic experiments, IP CNO or intra NAc CNO was administered to the mice prior to test start (>1 hr). For optogenetic experiments, mice were habituated to cable attachment prior to testing (>15 min). 3-point animal tracking was used to measure preference for scent with Ethovision 15 (Noldus, US) software.

#### Palatable food consumption

Single housed mice were habituated to ∼1g of Cocoa Pebbles cereal (Post, USA) in their home cage for 3 days prior to testing, also conducted in their home cage. Pre-weighed Cocoa Pebbles (∼1g) were placed in their cage, and intake was measured following 1hr (chemogenetic manipulation) or 10min (optogenetic manipulation). For chemogenetic experiments, IP CNO or intra NAc CNO was administered to the mice prior to start (>1 hr). For optogenetic experiments, the mice were habituated to cable attachment prior to test start (>15 min).

#### Open Field

During the active phase, and in low light (<15 lux measured on arena floor), mice were placed in one corner of an open field arena (L43 x W43 x H35 cm) and given free access to explore for 10 minutes. Animal movement was detected with an automated 3-point software (Ethovision). For analysis, the arena was subdivided into sixteen equal zones (4 inner and 12 outer squares). Ethovision 15 (Noldus, US) software calculated the time spent in each of the 16 zones. To measure time spent in center, the 4 inner zones were collapsed together.

#### Swim stress

Mice were placed in a plexiglass beakers (20 cm in diameter and 40 cm high) containing water (25ºC) filled to a depth of 30 cm, in which they could not touch the bottom of the beakers. The swim test lasted a maximum of 6 minutes. Behavior was captured using a video camera, and experimenters monitored the animal’s well-being throughout. Water was replaced and containers cleaned between individual mice. The durations of floating (immobility), climbing and swimming were scored manually for the final 4 minutes of recording. After testing, mice were towel dried and placed in a prewarmed cage before being returned to their home cage.

### Immunohistochemistry

To prepare tissue for the CRH staining, three CRH-Cre mice that had been injected with the Cre-driven AAV retrograde tracer in the NAc also received a single intracerebroventricular injection of colchicine (Tocris Bioscience, Bristol, UK, 10 μg/1 μl saline) 48 hours prior to perfusion to enhance CRH peptide localization in neuronal cell bodies. Coronal sections (20 μm) were subjected to CRH immunostaining. Briefly, after several washes with 0.01M PBS containing 0.3% Triton X-100 (PBS-T), sections were treated for 30 min in 0.3% H_2_O_2_-PBS-T, followed by blockade of nonspecific sites with 5% normal goat serum in PBS-T for 30 min. After rinsing with PBS three times, 5 minutes per wash, on a shaker at 95 rpm, sections were incubated for 48 hours at 4°C with rabbit anti-CRH primary antibody (courtesy of Paul E. Sawchenko, Salk Institute, PBL#rC68) in the blocking solution (dilution factor 1:10,000). The sections were rinsed with PBS-T three times, 5 minutes per wash, followed by incubation in an Alexa Fluor (AF) 488-conjugated goat-anti-rabbit secondary antibodies in the blocking solution (dilution 1:200, ThermoFisher, A-11008) for 2 hours. All the sections were rinsed with PBS-T three times, 5 minutes each. Sections were counter-stained with 10 μM DAPI (Sigma, D-9542), then wet-mounted on microscope slides and cover-slipped with mounting medium (Vectashield).

To assess the fidelity of CRH and the reporter, concurrent visualization of CRH and tdTomato was performed using the tyramide signal amplification (TSA) technique^47^. Sections (20 μm) were treated in 0.3% H_2_O_2_/PBS-T for 30 min, rinsed in PBS-T for 30 minutes and then blocked with 0.5% blocking buffer (PerkinElmer, FP120) for 2 hours. After rinsing of 10 minutes in PBS-T, sections were incubated with CRH rabbit antiserum (1:20,000) for 3 days (4°C), and then incubated in horseradish peroxidase (HRP)- conjugated anti-rabbit IgG (1:1,000; Perkin Elmer, Boston, MA) for 1.5 hours. Fluorescein-conjugated tyramide was diluted (1:150) in amplification buffer (Perkin Elmer, Boston, MA), and was applied in the dark for 5–6 min on ice.

### Image acquisition

Virus labeled sections were scanned under a 10x objective of a fluorescent microscope (Olympus BX 61) equipped with a high-sensitivity CCD camera (Nikon Eclipse E400). Confocal microscopy was utilized for imaging virus-labeled neurons after anterograde tracing (LSM 700, Zeiss). Images were obtained using Metamorph image acquisition software (Molecular Devices, Sunnyvale, CA) and analyses were done using Adobe Photoshop (CS4).

### Statistical analysis

All experiments and data analyses were conducted blind to experimental groups. The number of replicates (*n*) is indicated in figure legends and refers to the total number of experimental subjects independently treated in each experimental condition. Data are presented are mean values, and where applicable, accompanied by SEM. Statistical comparisons were performed using Prism 9 software (GraphPad, USA). Unpaired *t*-tests, paired *t*-tests, unpaired *u*-tests, and two-way ANOVA with repeated measures were used to test for statistical significance when appropriate. No statistical methods were used to pre-determine sample sizes. Statistical significance was set at α = 0.05. (NS, *P* > 0.05; \**P* < 0.05; \*\**P* < 0.01; \*\*\**P* < 0.001). *P* values are provided in all figure legends. Experiments were replicated in at least two independent batches, and from at least four different litters that yielded consistent results.

## Acknowledgements

This work was supported by National Institute of Health grants P50 MH096889, MH73136 and NS108296 (to T.Z.B) and the George E. Hewitt Foundation for Biomedical Research and the British Society for Neuroendocrinology (to M.T.B).

## Author contributions

M.T.B, A.K.S, B.G.G, L.Y.C, S.V.M, and T.Z.B contributed to study design. M.T.B, A.K.S, G.B.d.C and Y.C contributed to data collection. M.T.B and Y.C conducted all surgeries. M.T.B and A.K.S conducted all behavior studies. M.T.B, Y.C. and A.L.P conducted histological analyses. C.A.I and X.X provided anterograde and retrograde viruses. M.T.B and T.Z.B wrote the paper. All authors discussed and commented on the manuscript.

## Ethics declarations

The authors declare no competing interests.

## REFERENCES

1. Nestler, E. J. & Carlezon, W. A. The Mesolimbic Dopamine Reward Circuit in Depression. Biological Psychiatry vol. 59 1151–1159 (2006).

2. Koob, G. F. & Le Moal, M. Plasticity of reward neurocircuitry and the ‘dark side’ of drug addiction. Nature Neuroscience vol. 8 1442–1444 (2005).

3. Lüthi, A. & Lüscher, C. Pathological circuit function underlying addiction and anxiety disorders. Nat. Neurosci. 17, 1635–1643 (2014).

4. Nelson, C. A. et al. Adversity in childhood is linked to mental and physical health throughout life. BMJ 371, m3048 (2020).

5. Baldwin, J. R. et al. Population vs Individual Prediction of Poor Health From Results of Adverse Childhood Experiences Screening. JAMA Pediatr. 175, 385–393 (2021).

6. Berridge, K. C. & Kringelbach, M. L. Pleasure Systems in the Brain. Neuron vol. 86 646–664 (2015).

7. Yang, H. et al. Nucleus Accumbens Subnuclei Regulate Motivated Behavior via Direct Inhibition and Disinhibition of VTA Dopamine Subpopulations. Neuron 97, 434–449.e4 (2018).

8. Britt, J. P. et al. Synaptic and Behavioral Profile of Multiple Glutamatergic Inputs to the Nucleus Accumbens. Neuron 76, 790–803 (2012).

9. Beyeler, A. et al. Organization of Valence-Encoding and Projection-Defined Neurons in the Basolateral Amygdala. Cell Rep. 22, 905–918 (2018).

10. Ambroggi, F., Ishikawa, A., Fields, H. L. & Nicola, S. M. Basolateral Amygdala Neurons Facilitate Reward-Seeking Behavior by Exciting Nucleus Accumbens Neurons. Neuron 59, 648–661 (2008).

11. Stuber, G. D. et al. Excitatory transmission from the amygdala to nucleus accumbens facilitates reward seeking. Nature 475, 377–380 (2011).

12. Wassum, K. & Izquierdo, A. The basolateral amygdala in reward learning and addiction. Neurosci. Biobehav. Rev. 57, 271–283 (2015).

13. Peña, C. J. et al. Early life stress confers lifelong stress susceptibility in mice via ventral tegmental area OTX2. Science. 356, 1185–1188 (2017).

14. Birn, R. M., Roeber, B. J. & Pollak, S. D. Early childhood stress exposure, reward pathways, and adult decision making. Proc. Natl. Acad. Sci. 114, 13549–13554 (2017).

15. Birnie, M. T. & Baram, T. Z. Principles of emotional brain circuit maturation. Science. 376, 1055–1056 (2022).

16. Callaghan, B. L. & Tottenham, N. The Neuro-Environmental Loop of Plasticity: A Cross-Species Analysis of Parental Effects on Emotion Circuitry Development Following Typical and Adverse Caregiving. Neuropsychopharmacology 41, 163–176 (2016).

17. Luby, J. L., Baram, T. Z., Rogers, C. E. & Barch, D. M. Neurodevelopmental Optimization after Early-Life Adversity: Cross-Species Studies to Elucidate Sensitive Periods and Brain Mechanisms to Inform Early Intervention. Trends Neurosci. 43, 744–751 (2020).

18. Green, J. G. et al. Childhood Adversities and Adult Psychiatric Disorders in the National Comorbidity Survey Replication I. Arch. Gen. Psychiatry 67, 113 (2010).

19. Krugers, H. J. et al. Early life adversity: Lasting consequences for emotional learning. Neurobiol. Stress 6, 14–21 (2017).

20. Deussing, J. M. & Chen, A. The Corticotropin-Releasing Factor Family: Physiology of the Stress Response. Physiol. Rev. 98, 2225–2286 (2018).

21. Lemos, J. C. et al. Severe stress switches CRF action in the nucleus accumbens from appetitive to aversive. Nature 490, 402–6 (2012).

22. Baumgartner, H. M., Schulkin, J. & Berridge, K. C. Activating Corticotropin-Releasing Factor Systems in the Nucleus Accumbens, Amygdala, and Bed Nucleus of Stria Terminalis: Incentive Motivation or Aversive Motivation? Biol. Psychiatry 89, 1162–1175 (2021).

23. Chen, Y. & Baram, T. Z. Toward Understanding How Early-Life Stress Reprograms Cognitive and Emotional Brain Networks. Neuropsychopharmacology 41, 197–206 (2016).

24. Walsh, J. J. et al. Stress and CRF gate neural activation of BDNF in the mesolimbic reward pathway. Nat. Neurosci. 17, 27–29 (2014).

25. Dedic, N. et al. Chronic CRH depletion from GABAergic, long-range projection neurons in the extended amygdala reduces dopamine release and increases anxiety. Nat. Neurosci. 21, 803–807 (2018).

26. Tervo, D. G. R. et al. A Designer AAV Variant Permits Efficient Retrograde Access to Projection Neurons. Neuron 92, 372–382 (2016).

27. Itoga, C. A. et al. New viral-genetic mapping uncovers an enrichment of corticotropin-releasing hormone-expressing neuronal inputs to the nucleus accumbens from stress-related brain regions. J. Comp. Neurol. 527, 2474–2487 (2019).

28. Sharp, B. M. Basolateral amygdala and stress-induced hyperexcitability affect motivated behaviors and addiction. Transl. Psychiatry 7, e1194–e1194 (2017).

29. Mahler, S. V et al. Designer receptors show role for ventral pallidum input to ventral tegmental area in cocaine seeking. Nat. Neurosci. 17, 577–585 (2014).

30. Mahler, S. V. et al. Chemogenetic manipulations of ventral tegmental area dopamine neurons reveal multifaceted roles in cocaine abuse. J. Neurosci. 39, 503–518 (2019).

31. Malkesman, O. et al. The Female Urine Sniffing Test: A Novel Approach for Assessing Reward-Seeking Behavior in Rodents. Biol. Psychiatry 67, 864–871 (2010).

32. Chen, Y. et al. Converging, synergistic actions of multiple stress hormones mediate enduring memory impairments after acute simultaneous stresses. J. Neurosci. 36, 11295–11307 (2016).

33. Birnie, M. T. et al. Plasticity of the Reward Circuitry After Early-Life Adversity: Mechanisms and Significance. Biological Psychiatry vol. 87 875–884 (2020).

34. Peña, C. J. et al. Early life stress alters transcriptomic patterning across reward circuitry in male and female mice. Nat. Commun. 10, 1–13 (2019).

35. Levis, S. C. et al. Enduring disruption of reward and stress circuit activities by early-life adversity in male rats. Transl. Psychiatry 12, 251 (2022).

36. Molet, J. et al. Fragmentation and high entropy of neonatal experience predict adolescent emotional outcome. Transl. Psychiatry 6, e702–e702 (2016).

37. Walker, C. D. et al. Chronic early life stress induced by limited bedding and nesting (LBN) material in rodents: critical considerations of methodology, outcomes and translational potential. Stress vol. 20 421–448 (2017).

38. Malter Cohen, M. et al. Early-life stress has persistent effects on amygdala function and development in mice and humans. Proc. Natl. Acad. Sci. 110, 18274–18278 (2013).

39. Dennison, M. J. et al. Differential Associations of Distinct Forms of Childhood Adversity With Neurobehavioral Measures of Reward Processing: A Developmental Pathway to Depression. Child Dev. 90, (2019).

40. Volkow, N. D. et al. Addiction: Decreased reward sensitivity and increased expectation sensitivity conspire to overwhelm the brain’s control circuit. BioEssays 32, 748–755 (2010).

41. Whitton, A. E. et al. Blunted Neural Responses to Reward in Remitted Major Depression: A High-Density Event-Related Potential Study. Biol. Psychiatry Cogn. Neurosci. Neuroimaging 1, 87–95 (2016).

42. Gunn, B. et al. The endogenous stress hormone CRH modulates excitatory transmission and network physiology in hippocampus. Cereb. Cortex 27, 4182–4192 (2017).

43. Roberto, M. et al. Corticotropin Releasing Factor–Induced Amygdala Gamma-Aminobutyric Acid Release Plays a Key Role in Alcohol Dependence. Biol. Psychiatry 67, 831–839 (2010).

44. Jo, Y. S., Namboodiri, V. M. K., Stuber, G. D. & Zweifel, L. S. Persistent activation of central amygdala CRF neurons helps drive the immediate fear extinction deficit. Nat. Commun. 11, 422 (2020).

45. McCullough, K. M. et al. Genome-wide translational profiling of amygdala Crh-expressing neurons reveals role for CREB in fear extinction learning. Nat. Commun. 11, 5180 (2020).

46. de Guglielmo, G. et al. Inactivation of a CRF-dependent amygdalofugal pathway reverses addiction-like behaviors in alcohol-dependent rats. Nat. Commun. 10, 1238 (2019).

47. Chen, Y., Molet, J., Gunn, B. G., Ressler, K. & Baram, T. Z. Diversity of Reporter Expression Patterns in Transgenic Mouse Lines Targeting Corticotropin-Releasing Hormone-Expressing Neurons. Endocrinology 156, 4769–4780 (2015).

